# Colorectal cancer detection and treatment with engineered probiotics

**DOI:** 10.1101/2023.04.03.535370

**Authors:** Candice R. Gurbatri, Georgette Radford, Laura Vrbanac, Courtney Coker, Jong-won Im, Samuel R. Taylor, YoungUk Jang, Ayelet Sivan, Kyu Rhee, Anas A. Saleh, Tiffany Chien, Fereshteh Zandkarimi, Ioana Lia, Tamsin RM Lannagan, Tongtong Wang, Josephine A Wright, Elaine Thomas, Hiroki Kobayashi, Jia Q Ng, Matt Lawrence, Tarik Sammour, Michelle Thomas, Mark Lewis, Lito Papanicolas, Joanne Perry, Tracy Fitzsimmons, Patricia Kaazan, Amanda Lim, Julie Marker, Cheri Ostroff, Geraint Rogers, Nicholas Arpaia, Daniel L Worthley, Susan L Woods, Tal Danino

## Abstract

Bioengineered probiotics enable new opportunities to improve colorectal cancer (CRC) screening, prevention and treatment strategies. Here, we demonstrate the phenomenon of selective, long-term colonization of colorectal adenomas after oral delivery of probiotic *E. coli* Nissle 1917 (EcN) to a genetically-engineered murine model of CRC predisposition. We show that, after oral administration, adenomas can be monitored over time by recovering EcN from stool. We also demonstrate specific colonization of EcN to solitary neoplastic lesions in an orthotopic murine model of CRC. We then exploit this neoplasia-homing property of EcN to develop early CRC intervention strategies. To detect lesions, we engineer EcN to produce a small molecule, salicylate, and demonstrate that oral delivery of this strain results in significantly increased levels of salicylate in the urine of adenoma-bearing mice, in comparison to healthy controls. We also assess EcN engineered to locally release immunotherapeutics at the neoplastic site. Oral delivery to mice bearing adenomas, reduced adenoma burden by ∼50%, with notable differences in the spatial distribution of T cell populations within diseased and healthy intestinal tissue, suggesting local induction of robust anti-tumor immunity. Together, these results support the use of EcN as an orally-delivered platform to detect disease and treat CRC through its production of screening and therapeutic molecules.

## INTRODUCTION

Synthetic biology enables the engineering of microbes as living diagnostics and medicines through colonization of niches such as the gastrointestinal tract, skin, lung and tumors^1-3^. To date, a multitude of studies have shown that bacteria selectively colonize a broad range of tumor types, taking advantage of tumor hallmarks such as a necrotic core, hypoxia and reduced immune surveillance within the tumor microenvironment (TME). Here bacteria can be engineered to produce a range of payloads within the tumor, but studies thus far have mainly focused on intratumoral or systemic administration in subcutaneous tumor models^4-6^, leading to concerns over toxicity and translational relevance. Moreover, colonization of these delivery routes has been limited to milli-meter-size lesions, making early intervention challenging. Oral delivery of probiotics is a preferred method of administration and enables access to the gastrointestinal tract for local delivery of therapeutic or diagnostic molecules that can subsequently be non-invasively monitored. Recent microbial gene circuits have reported the ability of engineered probiotics to sense, record and respond to signals of inflammation and infection within the gut^7-11^. However, these approaches have relied on known biomarkers and have not yet detected intestinal cancers in the gut.

Colorectal cancer (CRC) is the second leading cause of cancer morbidity and mortality worldwide, with significantly rising incidence rates in younger populations, emphasizing the need for improved and affordable interventions^12^. Colonoscopy is effective at reducing CRC incidence and mortality, but is inconvenient, costly, and is associated with rare, but significant, complications^13, 14^. Furthermore, genetic conditions that can predispose patients to CRC, such as familial adenomatous polyposis (FAP), result in hundreds of colonic adenomas, the primary precursor lesions of CRC^13, 15-17^, complicating CRC prevention. Surgical interventions like polypectomy and colectomies are options for early-stage disease and can be used in combination with chemo(radio)therapies^18, 19^. Recently, favorable outcomes have been reported in clinical trials with anti-programmed cell death protein-1 (anti-PD1) checkpoint therapy in microsatellite-instability – high (MSI-H) CRC, but with limited success in microsatellite-stable (MSS) CRC disease, which represents ∼85% of CRC^20-22^. Thus, there is an unmet need for an approach that can target, detect, and treat adenomas to prevent progression to malignant disease.

Here, we describe the phenomenon of probiotic *E. coli* Nissle 1917 (EcN) to selectively colonize adenomas and isolated neoplastic lesions when orally delivered to orthotopic, genetic and transplant murine models of CRC. We demonstrate that these bacteria remain colonized long-term, and adenomas can be detected by recovering EcN from stool or by engineering EcN to produce a small molecule measurable in the urine. Lastly, we show that engineered EcN can deliver immunotherapeutics within adenomas, manipulating the TME *in situ* and reducing overall disease burden.

## RESULTS

CRC precursor lesions were modeled using *Apc*^*Min/+*^ mice^23^, a well-established mouse model of FAP, as they develop spontaneous intestinal adenomas and are representative of initiating genetic mutations seen in ∼80% of human CRC^24^ (**Fig. 1A**). We explored neoplasia colonization by orally-delivering EcN encoding a genomically-integrated *luxCDABE* cassette (EcN-lux)^25^ to *Apc*^*Min/+*^ mice with an intact immune system and microbiome. *In vivo* imaging of mice dosed with EcN-lux interestingly showed elimination of bioluminescent bacteria in healthy wild-type (WT) mouse gut, but retention in the *Apc*^*Min/+*^ mouse gut for up to 7 weeks after oral administration (**Fig. 1B**). Enrichment of EcN-lux in neoplasia was further demonstrated with *ex vivo* imaging of intestinal tissue, where bioluminescence co-localized with visible macroadenomas and generally, more bioluminescence was observed in the distal small intestine where polyp burden was the greatest (**Fig. 1C, Fig. S1**). In *Apc*^*Min/+*^ mice with and without orally delivered EcN, we did not observe any notable effects regarding mouse body condition and weight, as others have demonstrated previously with EcN strains in other mouse types^26-28^. Due to increased concerns of colibactin-producing bacteria like EcN being pro-carcinogenic^29^, we sought to disrupt colibactin production by deleting the *clbA* gene (EcN*ΔclbA*)^30, 31^ and investigated this engineered strain’s colonization ability (**Fig. S2A**). When delivered orally to tumor-bearing *Apc*^*Min/+*^mice, EcN*ΔclbA* survived transit through the gut as detectable in *Apc*^*Min/+*^ feces for multiple days after oral dosing. Furthermore, bioluminescent EcN*ΔclbA* co-localized with visible macroadenomas as observed by *ex vivo* intestinal imaging, suggesting that colonization does not rely on the presence of *clbA gene* or an intact colibactin-encoding operon (**Fig. S2B-C**). Subsequent plating of homogenized intestinal tissue on antibiotic-selective Luria broth (LB) plates specific for EcN-lux, indicated that no detectable EcN-lux could be recovered from wild-type mouse tissue, suggesting that a long-term niche is not formed in the gut unless neoplastic tissue is present (**Fig. 1D**). To further investigate bacteria localization, EcN-lux was engineered to produce a human influenza hemagglutinin (HA)-tagged reporter protein and orally delivered to *Apc*^*Min/+*^ mice. Immunohistochemical staining against HA in swiss-rolled intestinal tissue showed EcN-lux within adenomas of varying sizes, including those less than a millimiter (**Fig. 1E, Fig. S3**).

**Fig. 1.**
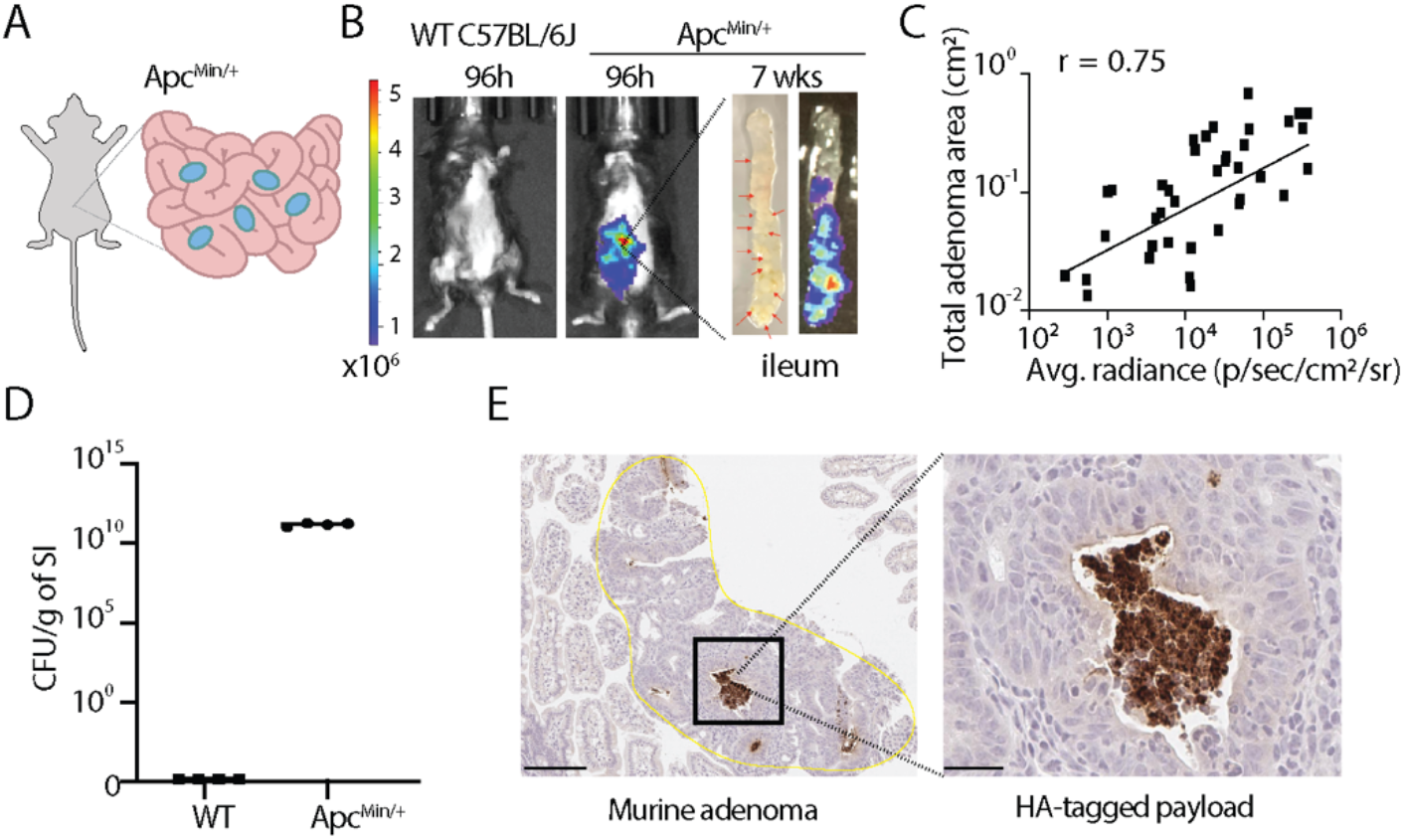
Adenoma colonization of *E. coli* Nissle 1917 (EcN) in a genetically-engineered mouse model of CRC. **(A)** Schematic of spontaneous intestinal adenomas in *Apc*^*Min/+*^ model. 12-week-old *Apc*^*Min/+*^ mice were gavaged twice, 3-4 days apart with EcN expressing luxCDABE luciferase cassette (EcN-lux). (**B)** EcN-lux visualized using an IVIS for bioluminescence *in vivo* 96h post dosing. After 7 weeks, mice were sacrificed, intestinal tissue was excised and *ex vivo* imaged for bioluminescence. Red arrows point to macroadenomas on distal intestinal tissue (n = 20 WT, n=25 *Apc*^*Min/+*^). **(C)** Plot where x-axis is bioluminescence signal measured from *ex vivo* images of bisected intestinal tissue and y-axis is total adenoma area in matched tissue sections as measured in H&E-stained images (r = 0.75, Spearman correlation coefficient). **(D-E)** In separate cohorts, mice were dosed with EcN-lux or EcN producing an HA-tagged protein and **(D)** sacrificed at 1 week post dosing and intestinal tissue was homogenized and plated on antibiotic-selective plates for EcN-lux to quantify colony-forming-units (CFU) per gram of tissue (n=5 WT, n=5 *Apc*^*Min/+*^) or **(E)** sacrificed at 4 weeks post dosing and intestinal tissue was paraffin-embedded, sectioned and stained by anti-HA immunohistochemistry. Dark brown stain depicts HA-tagged protein produced by EcN in adenomas.

We next tested selective colonization of isolated lesions by evaluating EcN-lux in an orthotopic model of CRC, representative of MSS adenocarcinomas in humans, whereby murine CRC organoids were injected into the distal murine colon and tumor grade tracked via weekly colonoscopy^32^ (**Fig. 2A, Fig. S4**). To home in on our specific bacteria of interest, mice were pre-treated with broad-spectrum antibiotics, which disrupts normal microbiota composition, a common phenomenon in gastrointestinal diseases including CRC^33^. EcN-lux was orally delivered and *in vivo* imaging five days post dosing revealed colocalization of bioluminescent EcN-lux with colon tumors (**Fig. 2B-C, S5**). Subsequent homogenization and plating of excised organs on antibiotic-selective LB plates confirmed EcN-lux was significantly enriched in tumors compared to adjacent healthy tissue and peripheral organs (**Fig. 2D**). The median diameter of EcN-lux colonized tumors was 2 mm (+/-1.2 mm), suggesting the size of neoplastic lesions detected using this EcN-lux platform was similar to colonoscopic reporting of diminutive (0–5 mm) polyps in humans^34^. Additional histological interrogation of these tumors also indicated the presence of tumor mucin lakes, demonstrating this model phenocopies mucinous human adenocarcinomas^32^(**Fig. 2E**), and that mucinous differentiation does not inhibit EcN-lux colonization. Indeed, specific localization of EcN-Lux in these tumors by RNA *in situ* hybridization, suggest that the bacteria can predominantly be found in nests on the luminal tumor surface and can co-locate with hypoxic tumor regions that may further facilitate bacterial growth within the tumor space (**Fig. 2F, S6**).

**Fig. 2.**
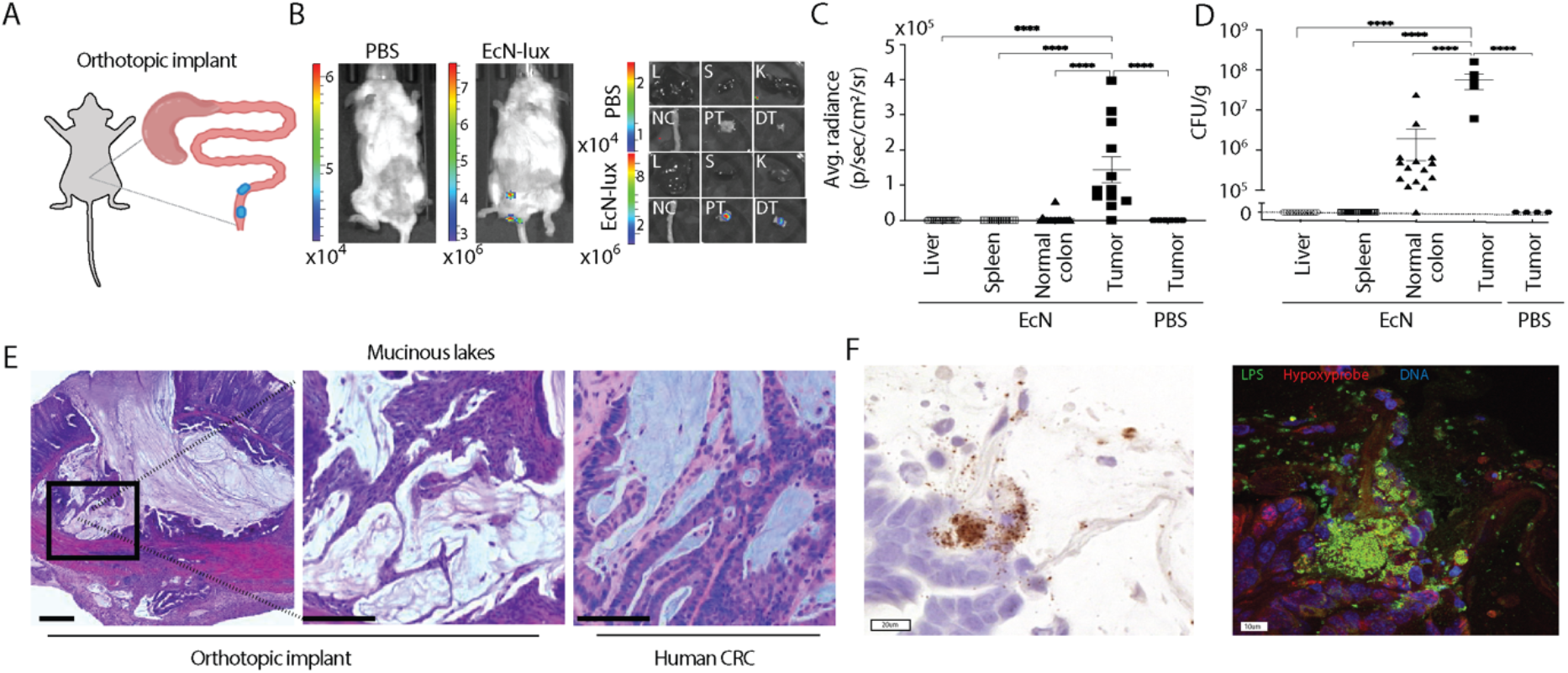
Tumor colonization of *E. coli* Nissle 1917 (EcN) in an orthotopic mouse model of of CRC. **(A)** Schematic of orthotopic, distal CRC model generated by mucosal injection of CRC organoids. Tumor growth was monitored by colonoscopy, with tumor and non-tumor bearing control animals orally-dosed twice, 2 days apart, with EcN-lux or PBS before **(B)** imaging 5 days after last dose for bioluminescence, with **(C)** luminescence quantified in organs *ex vivo* (L, liver n = 17, S, spleen n = 17, NC, normal colon n = 17, PT, proximal tumor, DT distal tumor, n = 12, PBS tumors n = 6, **** p < 0.0001, one-way ANOVA with Tukey’s multiple comparisons). **(D)** Tissues were homogenized, plated on antibiotic-selective plates, and quantified for CFU per gram (Liver n = 17, Spleen n = 17, Normal Colon n = 17 from EcN-dosed non-tumor mice and EcN-dosed tumor mice n = 6, PBS-dosed tumor mice n = 4). **** p < 0.0001, one-way ANOVA with Tukey’s multiple comparisons). **(E)** Histopathology of orthotopic tumor (left) and higher power image of boxed region (middle) showing tumor mucin lakes similar to (right) human CRC with overt mucous phenotype.(**F**) Representative images of orthotopic CRC from mouse orally-dosed twice, 2 days apart, with EcN-lux, with serial sections showing (left) EcN-lux specific location by RNAscope in situ hybridization for *lux* (brown, scale bar 20µm) and (right) immunofluorescent staining for Hypoxyprobe (red) and lipopolysaccharide (LPS, green, scale bar 10µm).

This phenomenon of neoplasia selective colonization of EcN suggested its utility as a platform to both monitor adenoma presence and also then to manipulate lesions *in situ*. Since stool-based tests are a common non-invasive screening tool available for CRC, we first determined if EcN-lux extracted from stool could be used to non-invasively monitor the presence of adenomas over time. To do this, we orally dosed both healthy wild-type (WT) and *Apc*^*Min/+*^ mice with EcN-lux, collected stool pellets at predetermined time points, then homogenized and plated fecal matter on antibiotic-selective LB agar plates (**Fig. 3A**). During the first seven hours post dosing, both WT and *Apc*^*Min/+*^ mice had comparable shedding of EcN-lux CFU in their stool, corresponding to material transit time through the gut^35^. However, by twenty-four hours, levels of EcN-lux were undetectable in some healthy mice and by forty-eight hours we were unable to recover EcN-lux from the stool of dosed WT mice (**Fig. 3B**). Furthermore, EcN-lux maintained its genomically-encoded luminescence cassette, allowing for simple luminescence imaging to screen for EcN-lux positive colonies as a proxy for adenoma presence over time. While this stool test could be a useful method for adenoma tracking, we aimed to investigate the more clinically relevant utility of EcN as a diagnostic platform. To do this, we sought to have EcN produce a molecule that could be more conveniently recovered from bodily fluids and ultimately engineered the production of salicylate, which can be feasibly detected in urine (**Fig. 3D**)^36^. We thus engineered EcN-lux to express, *pchA* and *pchB*, enzymes critical to the shikimate pathway, present in bacteria, in converting chorismite to salicylate^37^. Liquid chromatography-mass spectrometry (LC-MS) was performed to confirm salicylate production in supernatant and cell pellet samples of EcN-SA, with a measurable concentration of ∼2uM per 1×10^8^ EcN-SA (**Fig. 3E, S7**). After quantifying the salicylate production of our engineered probiotic, we sought to evaluate its behavior *in vivo* and its potential to serve as a urine-based adenoma screening method. To do this, we first acquired a baseline urine sample and then dosed WT and *Apc*^*Min/+*^ mice orally with the EcN-SA strain and collected urine twenty-four hours later, when a majority of the probiotic would have transited out of the healthy WT gut, but when specificity was low with the stool-based method (**Fig. 3B**). We then probed for salicylate presence using LC-MS. While urine from both the WT and *Apc*^*Min/+*^ had detectable levels of salicylate, the tumor-bearing mice had approximately 20 times more relative salicylate in the urine (**Fig. 3F**), with absolute concentration ranging from 2-10uM. Taken together, these data suggest that engineered bacteria can be delivered to adenoma-bearing mice, maintain their genetic circuitry, and be used as a proxy for non-invasive adenoma tracking and possible early detection in both fecal matter and urine-based assays.

**Fig. 3.**
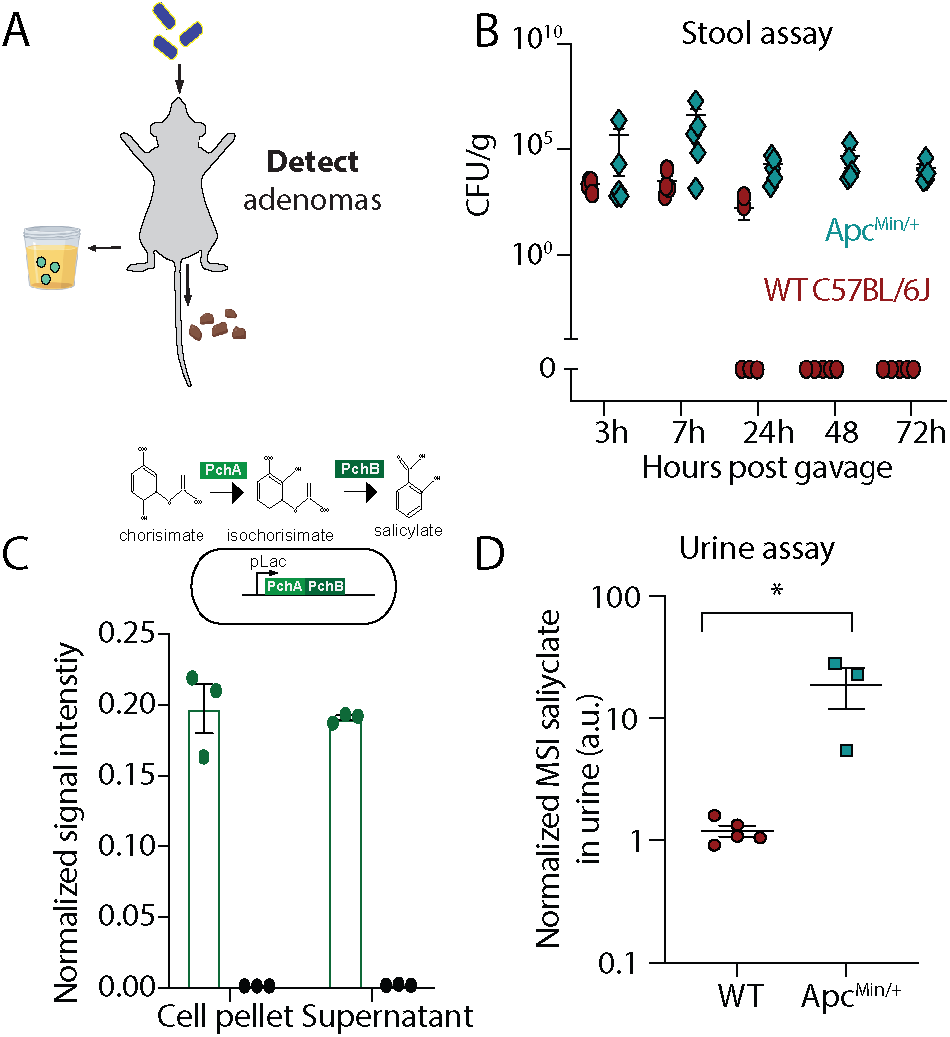
Engineering of stool and urine-based EcN platform for non-invasive adenoma tracking. **(A)** Schematic of orally-delivered EcN probiotic to be detected in fecal matter by quantifying colony-forming units. **(B)** 15-week-old *Apc*^*Min/+*^ mice were dosed orally 3-4 days apart with EcN and one stool pellet was collected at 3, 7, 24, 48, and 72h after last dose, homogenized, plated on antibiotic-selective plates and quantified for CFU (n=5 mice per group). (**C)** Schematic of the metabolic pathway whereby enzymes *pchA* and *pchB* convert chorismate to salicylate (Top). These enzymes were cloned onto plasmids and transformed into EcN (EcN-SA). Overnight cultures of EcN and EcN-SA were optical density-matched and LC-MS was used to detect salicylate in both the cell pellet and supernatant of EcN and EcN-SA cultures (Bottom). All samples were normalized to an internal isotope-labelled D4-salicylate standard. **(D)** 15-week-old *Apc*^*Min/+*^ mice were dosed with 10^9^ EcN-SA bacteria and urine was collected 24hr after dosing. LC-MS quantification of salicylate molecules in urine of wild-type (WT) and *Apc*^*Min/+*^ mice normalized to pretreated urine values (p<0.05, unpaired T test, n=3-5 mice per group).

With the ability to non-invasively determine adenoma presence, we next sought to address whether our screening system could be adapted for therapeutic purposes and reduce polyp burden. As the *Apc*^*Min/+*^ model is considered to be microsatellite stable (MSS), which traditionally responds poorly to immunotherapeutic approaches, we leveraged our probiotic delivery platform, whereby the bacteria serve as an immune adjuvant and also a chassis to deliver multiple immunotherapeutic payloads. Given that we have confirmed that EcN can maintain its genetic circuitry when delivered orally, we combined therapeutic delivery with an EcN-lux strain genomically-encoded with a lysis circuit optimized (SLIC) to maximize immunotherapeutic release^38, 39^. SLIC was used to deliver nanobodies blocking PD-L1 and CTLA-4 targets and cytokines like GM-CSF (SLIC-3), which has been shown to work in combination to enhance efficacy of checkpoint blockade therapy in a subcutaneous mouse colorectal model when delivered intratumorally^39^ (**Fig. 4A**). Here, the *Apc*^*Min/+*^ mice were either orally dosed twice with PBS or EcN-lux strains and then sacrificed ∼1 month later. Histological analysis of hematoxylin and eosin-stained tumors demonstrate an overall reduction of adenoma area (**Fig. 4B**) and number (**Fig. S8A-B**) by ∼47% with SLIC-3 treatment. Notably, an increased percentage of smaller adenomas was observed in SLIC-3 treated mice, whereas PBS-treated mice tended to have larger adenomas (**Fig. S8C**). Moreover, this reduction was not specific to a location and was observed throughout the small intestine (**Fig. 4C-D**). Interrogation of immunophenotype on tissue sections suggests that reduction in adenoma burden is associated with increased infiltration of CD3+, CD8+ cells within adenomas and increased production of intratumoral granzymeB, suggesting immune-mediated tumor cell killing in SLIC-3 treated mice (**Fig. 4E-G**).

**Fig. 4.**
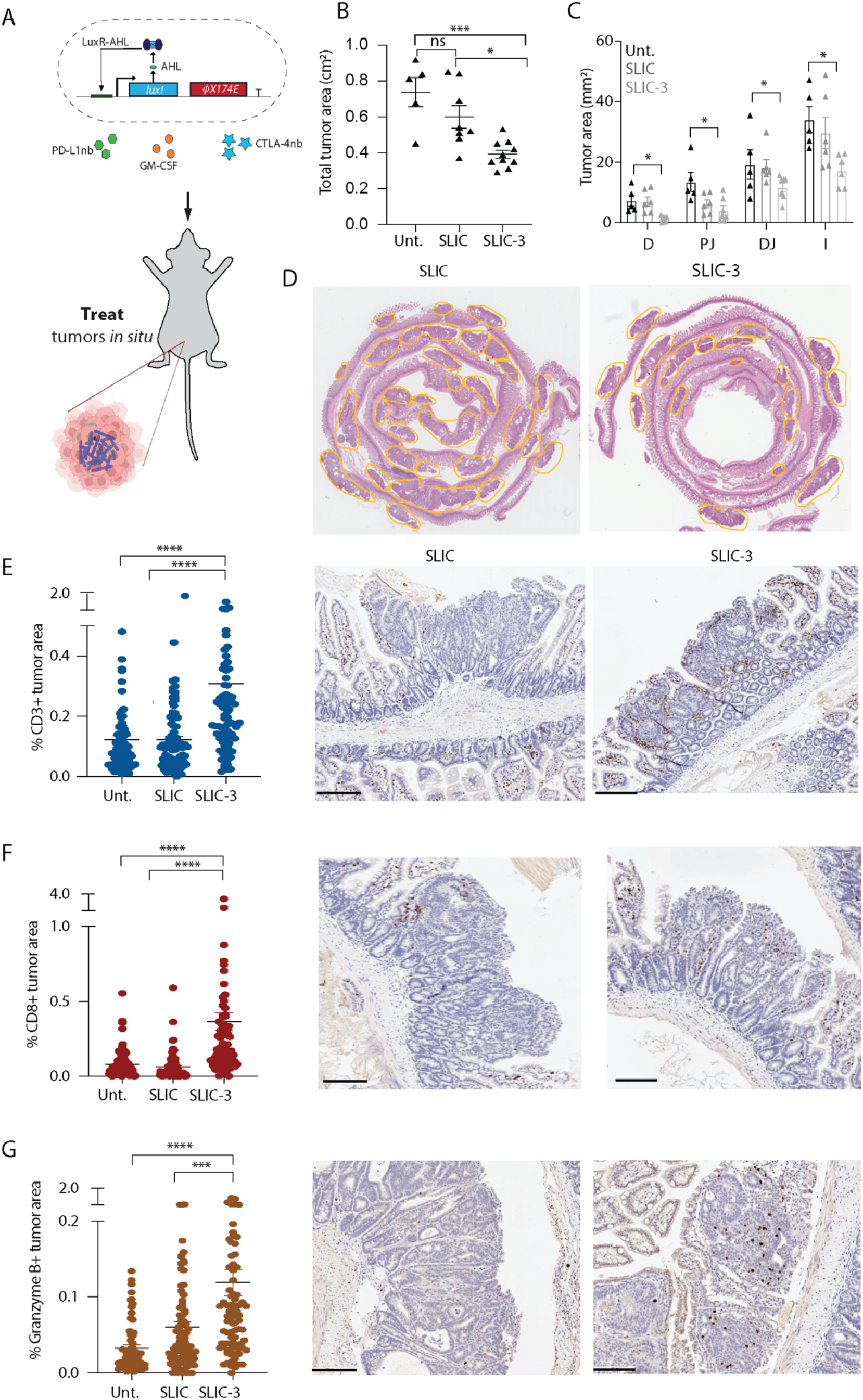
Treatment with EcN engineered to produce immunotherapeutics reduces adenoma burden and modifies the tumor-immune microenvironment. **(A)** Schematic of orally-delivered EcN probiotic engineered to lyse and produce immunotherapeutic proteins *in situ*. **(B-C)** 15-week-old *Apc*^*Min/+*^ mice were dosed with PBS (Unt), EcN genomically encoding a lysis circuit (SLIC) or SLIC producing granulocyte-macrophage colony-stimulating factor (GM-CSF), and blocking nanobodies against PD-L1 and CTLA-4 targets (SLIC-3). 1 month after dosing, mice were sacrificed, intestines were bisected, swiss-rolled, paraffin embedded, sectioned, stained with hemotoxin and eosin and quantified for **(B)** overall tumor area and **(C)** tumor area along the intestine, D, duodenum, PJ, proximal jejunum, DJ, distal jejunum, I, ileum (* p<0.05, ** p < 0.01, *** P, 0.001, ns, not significant, ordinary one-way ANOVA test with Holm-sidak multiple comparisons test, n=5-10 mice per group). **(D)** Representative H&E-stained histology images of SLIC and SLIC-3 treated mice. **(E-G)** Using immunohistochemical (IHC) techniques, intestinal tissue sections from all mouse groups were stained and quantified for **(E)** CD3+, **(F)** CD8+, and **(G)** granzymeB+ cells (n=3-5 mice per group, each dot represents a polyp, n=50-100 polyps per group, **** p < 0.0001 ordinary one-way ANOVA with Tukey post-test).) Representative IHC images of all three markers are shown beside their respective plots with positive staining depicted as brown puncta in SLIC and SLIC-3-treated mice. Scale bars represent 200 µm.

## DISCUSSION

Altogether, we have demonstrated selective and robust colonization of adenomas and tumors in two distinct orthotopic murine models, by orally-delivered probiotic EcN. Leveraging this colonization ability, we demonstrated the possibility for engineered EcN in adenoma diagnosis through non-invasive stool and urine assays. Furthermore, we demonstrated therapeutic potential by encoding EcN to produce checkpoint inhibitor therapies and cytokine GM-CSF, thereby significantly reducing adenoma burden in a model of MSS CRC through oral delivery, a disease subtype that in humans is normally unresponsive to systemically-administered checkpoint inhibitors^40, 41^. Moreover, additional benefits were observed in using a probiotic platform compared to conventional checkpoint therapies, including possible remodeling of the TME. Interestingly, T cells were detected at the periphery of polyps in the untreated group, consistent with multiple studies demonstrating a correlation between WNT/beta-catenin activation characteristic of lesions in *Apc*^*Min/+*^ mice and T cell exclusion^42, 43^. These data also demonstrate enhanced T cell infiltration into the polyp core upon bacterial-production of GM-CSF and checkpoint inhibitor nanobodies, potentially overcoming WNT/beta catenin-mediated T cell exclusion; however, more experimentation is necessary to understand the underlying mechanisms involved. Finally, since our probiotic platform is modular, there is the possibility to expand both screening and therapeutic cargo to explore other therapeutic combinations and achieve enhanced adenoma reduction.

While we explored colonization of EcN across multiple murine models both with intact and disrupted microbiomes, more investigation into the effects of the native tumor microbiome on EcN colonization could provide insight into generalizability of this approach across patients with varying diets and microbiota, including those who might already have baseline levels of EcN present^44^. Furthermore, a deeper understanding of characteristics governing EcN establishment within tumors is needed. The presented platform demonstrated that engineered probiotic strains maintain their programmed behavior within the complex gut environment, highlighting their potential for a range of diagnostic and therapeutic applications. Taken together, these data lay the groundwork for future pre-clinical and clinical experiments testing engineered EcN for CRC screening, prevention and treatment.

## MATERIALS AND METHODS

### Strains and plasmids

All bacterial strains used were luminescent (integrated *luxCDABE* cassette) so they could be visualized with the In Vivo Imaging System (IVIS)^45^. The EcN*ΔclbA* strain was engineered using the lambda-red recombineering method^46^. The salicylate-encoding plasmid was constructed using Gibson assembly methods or restriction enzyme-mediated cloning methods whereby *pchA* and *pchB* genes were cloned onto a high-copy origin plasmid and driven by the *lac* promoter. The SLIC and SLIC-3 strains were constructed as previously described^39^.

### Bacterial preparation for oral administration

Overnight cultures of EcN-lux were diluted 1:100 into LB with 50 ng/ml erythromycin and cultured to an OD600 of 0.1-0.5 on a shaker at 37 °C. Bacteria were collected by centrifugation at 3000-5000 r.c.f., washed three times with sterile PBS, resuspended in sterile ice-cold PBS with a total of 100-200uL dosed at orally at a concentration of ∼10^10^ CFU/ml. SLIC strains were prepared as previously described^39^. Briefly, growth media for SLIC and SLIC-3 strains also contained 0.2% glucose to suppress premature lysis in culture. Additionally, SLIC-3 strains were grown with 50 µg/ml kanamycin.

### Organoid culture

Mouse CRC *Braf*^*V600E*^*;Tgfbr2*^Δ*/*Δ^*;Rnf43* ^Δ*/*Δ^*/Znrf3* ^Δ*/*Δ^*;p16 Ink4a* ^Δ*/*Δ^ (*Braf*^*V600E*^Δ*TRZI)* organoids were generated using CRISPR/Cas9 genome engineering and expanded for injection into mice in matrigel culture as described^32^. Culture medium was Advanced Dulbecco’s modified Eagle medium/F12 (Life Technologies) supplemented with 1x gentamicin/antimycotic/antibiotic (Life Technologies), 10mM HEPES, 2mM GlutaMAX, 1xB27 (Life Technologies), 1xN2 (Life Technologies), 50ng/ml mouse recombinant EGF (Peprotech), 100 ng/ml mouse recombinant noggin (Peprotech), 10ng/ml human recombinant TGF-β1 (Peprotech). Immediately after each split, organoids were cultured in 10uM Y-27632 (In Vitro Technologies), 3uM iPSC (Calbiochem Cat #420220), 3uM GSK-3 inhibitor (XVI, Calbiochem, # 361559) for the first 3 days.

### Orthotopic mouse model of CRC

All animal experimentation related to the orthotopic CRC model was approved by the institutional animal ethics committee (SAM-319, SAM-20-031). Orthotopic injections to generate distal colon tumors were undertaken as previously described^47^. In brief, *NOD*.*Cg-Prkdc*^*scid*^*Il2rg*^*tm1Wjl*^*/SzJ* (NSG) mice (male and female,6–12 weeks old) were obtained from the SAHMRI Bioresources facility and housed under SPF conditions. Digested *Braf*^*V600E*^*ΔTRZI* organoid clusters (equivalent to ∼150 organoids) were resuspended in 20 μL 10% GFR matrigel 1:1000 india ink, 10 μM Y-27632 in PBS and injected into the mucosa of the distal colon of anaesthetised NSG mice using colonos-copy-guided orthotopic injection (2 injection sites/mouse). Injection sites were monitored by weekly colonoscopy. EcN administration began once the tumors were clearly established at grade 3 to 4^48^, 3 weeks post organoid injection. Broad-spectrum antibiotic treatment to generate gut dysbiosis in specific groups involved administration of 0.5g/L neomycin and 1g/L ampicillin in *ad libitum* drinking water for 5 days. This that was halted 6h prior to EcN administration.

### *Apc*^*Min/+*^ mouse model of CRC

All animal experimentation related to the *Apc*^*Min/+*^ mouse model of CRC was approved by the Institutional Animal Care and Use Committee (Columbia University, protocols AC-AAAN8002 and AC-AAAZ4470). All mice were regularly monitored and euthanized based on veterinarian recommendation or when they reached ∼20 weeks of age. In all therapeutic studies wild-type (WT) littermates of *Apc*^*Min/+*^ on the C57BL/6 background were used and both males and females were treated and evenly distribute among groups. For diagnostic studies, of *Apc*^*Min/+*^ mice were purchased from Jackson Laboratories and purchased C57BL/6 mice were used as WT.

### Biodistribution

#### IVIS imaging

Background (stage alone) subtracted total flux (photons/second) was used to capture the light signal emitted by EcN in identically sized areas for each live mouse *in vivo*. Following necropsy, individual tissues were collected into individual wells of a 6-well plate, weighed and average radiance (photons/s/cm^2/sr) used for ex vivo tissue imaging to correct for the area being measured which differed for each tissue analyzed.

#### CFU

Excised tissues were placed asceptically into 5ml 20% glycerol in PBS and homogenized in MACS Gentle cell dissociator C tubes, one tissue per tube using program C. 100ul of each tissue homogenate glycerol stock was serially diluted 1:100 six times. 10ul of each dilution was spotted onto an LB agar plate with erythromycin selection at 50 µg/ml with 5 technical replicates. Plates were incubated at 37 °C overnight (16 hours). Colony forming units (CFU) were calculated for each sample normalized to weight of tissue input to generate CFU/g tissue. To generate CFU/g stool, one pellet of stool was placed into an Eppendorf and manually homogenized in PBS with a pipette tip and rigourous pipetting. Serial dilutions were spotted onto an LB agar place with 50ug/ml erythromycin and incubated at 37 °C overnight. CFU was normalized to weight of the stool.

### Salicylate metabolite detection

#### In vitro sample preparation

Overnight cultures of EcN-SA were OD600 matched to be 1. Cultures were then centrifuged at 3000 r.c.f. and 1mL of the supernatant was collected and stored at - 80°C, the rest was decanted, and the pellet was stored at -80°C as well until analysis. To 1ml of supernatants, 1000µL of extraction solvent ¬(MeOH/MeCN/H2O (2:2:1; v/v/v) containing 0.1mg/mL of D4-salicylate) was added. Similarly, to cell pellets, 800uL of the same extraction solvent with the internal D4-salicylate standard was added. Samples were homogenized using a Bead Ruptor 4 at speed 4 for 10 seconds for 5 cycles. The homogenized samples were centrifuged for 5 min at 14,000 x g. Then, the supernatant was removed and dried on the Gene Vac for 5 hours and re-suspended in 200µL of MeCN/H2O be-fore LC-MS analysis. A quality control (QC) sample was prepared by combining 20μL of each sample to assess the reproducibility of the features through the runs.

#### Ultra-High Performance Liquid Chromatography (UPLC) analysis

Chromatographic separation was carried out at 40°C on Acquity UPLC BEH C18 column (2.1× 50 mm, 1.8 μm) over a 7-min gradient elution. Mobile phase A consisted of water and mobile phase B was acetonitrile both containing 0.1% formic acid. After injection, the gradient was held at 99% mobile phase A for 0.5 min. For the next 4 min, the gradient was ramped in a linear fashion to 50% B and held at this composition for 1 min. The eluent composition returned to the initial condition in 0.1 min, and the column was re-equilibrated for an additional 1 min before the next injection was conducted. The flow rate was set to 450 μL/min and Injection volumes were 2 μL using the flow-through needle mode in the negative ionization mode. The QC sample was injected between the samples and at the end of the run to monitor the performance and the stability of the MS platform.

#### Mass Spectrometry (MS) analysis

The Synapt G2-Si mass spectrometer (Waters, Manchester, U.K.) was operated in the negative electrospray ionization (ESI) modes. A capillary voltage of - 1.5 kV and a cone voltage of 30 V was used. The source temperature was 120°C, and desolvation gas flow was set to 850 L/hr. Leucine enkephalin was introduced to the lock mass at a concentration of 2 ng/μL (50% ACN containing 0.1% formic acid), and a flow rate of 10 μL/min for mass accuracy and reproducibility. The data was collected in duplicates in the centroid data-independent (MSE) mode over the mass range m/z 50 to 650 Da with an acquisition time of 0.1 seconds per scan.The QC sample, D4-salicylate, and salicylate standards were also acquired in enhanced data-independent ion mobility (IMS-MSE) in negative modes for the structural assignment. The ESI source settings were the same as described above. The traveling wave velocity was set to 650 m/s and the wave height was 40 V. The helium gas flow in the helium cell region of the ion-mobility spectrometry (IMS) cell was set to 180 mL/min to reduce the internal energy of the ions and minimize fragmentation. Nitrogen as the drift gas was held at a flow rate of 90 mL/min in the IMS cell. The low collision energy was set to 4 eV, and the high collision energy was ramped from 25 to 50 eV in the transfer region of the T-Wave device to induce fragmentation of mobility-separated precursor ions.

#### Data pre-processing and statistical analysis

All raw data files were converted to netCDF format using DataBridge tool implemented in MassLynx software (Waters, version 4.1). Then, they were subjected to peak-picking, retention time alignment, and grouping using XCMS package (version 3.2.0) in R (version 3.5.1) environment. Technical variations such as noise were assessed and removed from extracted features’ list based on the ratios of average relative signal intensities of the blanks to QC samples (blank/QC >1.5). Also, peaks with variations larger than 30% in QCs were eliminated. The detected signal intensity of salicylate in the samples were normalized to the signal intensity of labeled D4-salicylate. Group differences in measured salicylate levels were calculated using the Welch t-test, p-value < 0.05 in GraphPad prism.

#### Urine sample preparation

Urine samples were collected from mice 24hours after EcN-SA oral dosing and frozen at -80°C for later LC-MS analysis. For LC-MS analysis, urine samples were thawed on ice and polar metabolites were extracted with addition of 100% ice-cold LC-MS grade methanol with 0.1M formic acid (4:1 ratio of extraction solvent volume to urine). Samples were then vortexed and centrifuged at 20,000g for 10 minutes at 4°C. Supernatants were then transferred to clean LC-MS tubes and loaded onto the LC-MS autosampler, which was temperature-controlled at 4°C.

#### Liquid Chromatography – Mass Spectrometry analysis

Targeted analysis was performed as previously described^49^. In brief, samples were separated by liquid chromatography on an Agilent 1290 Infinity LC system by injection of 3μl of extract through an Agilent InfinityLab Poroshell 120 HILIC-Z, 2.1 × 150 mm, 2.7μm (Agilent Technologies) column heated to 50 °C. Solvent A (100% water containing 10 mM ammonium acetate, 5 mM InfinityLab Deactivator Additive and adjusted to pH 9 using ammonium hydroxide) and Solvent B (85% acetonitrile/15% water containing 10 mM ammonium acetate, 5mM InfinityLab Deactivator Additive and adjusted to pH 9 using ammonium hydroxide) were infused at a flow rate of 0.250 ml min^−1^. The 26-min normal phase gradient was as follows: 0–2 min, 96% B; 5.5–8.5 min, 88% B; 9–14 min, 86% B; 17 min, 82% B; 23–24 min, 65% B; 24.5–26 min, 96% B; followed by a 10-min post-run at 96% B. Acquisition was performed on an Agilent 6230 TOF mass spectrometer (Agilent Technologies) using an Agilent Jet Stream electrospray ionization source (Agilent Technologies) operated at 3,500 V Cap and 0 V nozzle voltage in extended dynamic range, negative mode. The following settings were used for acquisition: The sample nebulizer set to 35 psi with sheath gas flow of 12 L min^−1^ at 350 °C. Drying gas was kept at 350 °C at 13 L min^−1^. Fragmentor was set to 90 V, with the skimmer set to 45 V and Octopole Vpp at 750 V. Samples were acquired in centroid mode at 1 spectra/s for *m/z* values from 50-1,700.

#### Data acquisition and analysis

Raw data was acquired from the instrument and analyzed using previously described open-source XCMS software^50^. Metabolites were identified from (*m/z*, rt) pairs by retention time comparison with authentic standards.

### Histology

For the *Apc*^*Min/+*^ mouse model, all intestinal tissue was excised with the caecum removed and tissue was bisected such that there were 5 total sections: duodenum, proximal jejunum, distal jejunum, ileum, and colon. Intestines were flushed with PBS, splayed open, swiss-rolled, and fixed overnight in 4% paraformaldaheyde. After 24h the swiss rolls were switched to 70% ethanol and sent for histology services at Histowiz, where they were paraffin-embedded, sectioned, and stained with either H&E or specific immunohistochemistry markers (HA-Tag C29F4 #3724 from Cell Signaling Technology; GranzymeB Leica Biosystems PA0291; CD3 Abcam 16669; CD8 catalog #CST98941 clone D4W2Z). Tumor sizes and IHC quantification were determined using FIJI software image analysis tools.

For the orthotopic CRC transplant model, mice were intraperitoneally injected with Pimonidazole-HCl (Hypoxyprobe, 60mg/kg) or PBS negative control for Hypoxyprobe staining, 1 hour prior to euthanasia. Tumor, nearby adjacent colonic tissue and kidneys were collected at necropsy, rinsed in PBS and fixed overnight in formalin prior to dehydration in 70% ethanol and paraffin-embedding. An additional positive control tumor sample for optimizing ISH staining was generated by intratumoral injection of EcN-lux into a dissected tumour sample from a PBS treated mouse *ex vivo*, prior to fixation. Formalin-fixed and paraffin-embedded tissue sections were stained with H&E. Serial sections were also subjected to co-immunofluorescence against Hypoxyprobe (cat# HP12-200 Kit, 1: 200 dilution) and lipopolysaccharide (LPS, cat# HM6011-100UG, 1:500 dilution) or chromogenic *in situ* hybridization using RNAscope technology (RNAscope 2.5 Detection Kit, Advanced Cell Diagnostics) following the manufacturer’s instructions with a custom probe to detect the *lux* transcript in EcN-lux or negative control probe, *DapB* (target region 414-862; catalogue number 310043). Briefly for ISH, tissue sections were baked in a dry oven (HybEZ II Hybridization System, ACD) at 60°C for 1 h and deparaffinized, followed by incubation with Hydrogen Peroxide (Lot# 322000, ACD) and targeted retrieval (Lot# 322330, ACD). Slides were incubated with relevant probes for 2 h at 40°C, followed by successive incubations with Amp1 to 6 reagents. Staining was visualized with DAB. For IF studies, sections were treated with blocking buffer (X0909, Dako) for 30 min, incubated with the indicated primary antibodies overnight at 4 °C, and washed with PBS. Sections were then incubated with Alexa Fluor 488/594-conjugated secondary antibodies (1:200 dilution, Thermo Fisher Scientific) for 1 h at room temperature. The sections were then mounted with Vectashield antifade mounting medium (Cat# H-1000-10, Vector Laboratories), and fluorescence was examined using a confocal laser-scanning microscope (FV3000, Olympus).

## Author contributions

C.R.G and C.C. performed the *in vivo* colonization studies in the *Apc*^*Min/+*^ model, G.R., T.R.M.L., T.W. in the orthotopic CRC model. C.R.G. analyzed the *Apc*^*Min/+*^ histology images with input and guidance from S.R.T. H.K. provided pathology analysis of orthotopic CRC model. C.R.G., S.R.T. and G.R. performed the IHC and ISH staining not performed by Histowiz. J.I, Y.U, T.C, designed and cloned the salicylate-producing strain. C.R.G., J.I., and Y.U., designed and/or engineered the EcNΔ*clbA* strains. C.R.G. performed all *ex vivo* assays in the *Apc*^*Min/+*^ model with assistance from I.L and C.C. G.R. performed all *ex vivo* assays in the orthotopic CRC model with assistance from L.V., J.A.W., and E.T. F.Z. performed all cell-culture-based mass spectrometry and A.A.S and K.R. performed all urine-based mass spectrometry. T.W., E.T., J.Q.N., monitored animals for orthotopic CRC model. J.M., C.O., G.R., M.L., T.S., M.T., M.L., L.P., J.P., T.F., P.K., A.L., T.D., D.L.W., C.R.G. S.L.W., provided intellectual input into study design. C.R.G., A.S., T.D., S.L.W., and D.L.W., wrote the manuscript with input from all authors.

