## Supplementary_Materials for "Colorectal cancer detection and treatment with engineered probiotics"

**List of Supplementary Materials**

Fig. S1. Orally-delivered EcN colonizes intestinal adenomas

Fig. S2. Colibactin is not a requisite for EcN colonization of intestinal adenomas

Fig. S3. EcN can colonize and release payloads within adenomas of varying stage

Fig. S4. Establishment of orthotopic CRC mouse model

Fig. S5. Schematic of experimental timeline

Fig. S6. Staining for the presence of EcN in orthotopic tissues

Fig. S7. EcN can produce salicylate molecules to be detected by liquid-chromatography mass spectrometry

Fig. S8. Orally-delivered EcN producing PD-L1 and CTLA-4 blocking nanobodies and GM-CSF reduces tumor burden in *Apc*$^{Min/+}$ mice

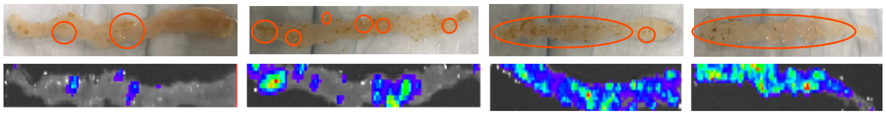


**Fig. S1. Orally-delivered EcN colonizes intestinal adenomas.** 12-week-old *Apc*$^{Min/+}$ mice were gavaged twice, 3-4 days apart with $10^9$ CFU bioluminescent EcN (EcN-lux). After 7 weeks, mice were sacrificed, intestinal tissue was excised and *ex vivo* imaged for bioluminescence. Red circles (top images) indicate areas of macroadenomas on sections of intestinal tissue isolated from the duodenum, proximal jejunum, distal jejunum and ileum. Bottom images show EcN-lux on the intestinal tissue sections. Figure S1 corresponds to data shown in **Fig. 1 B-C.**

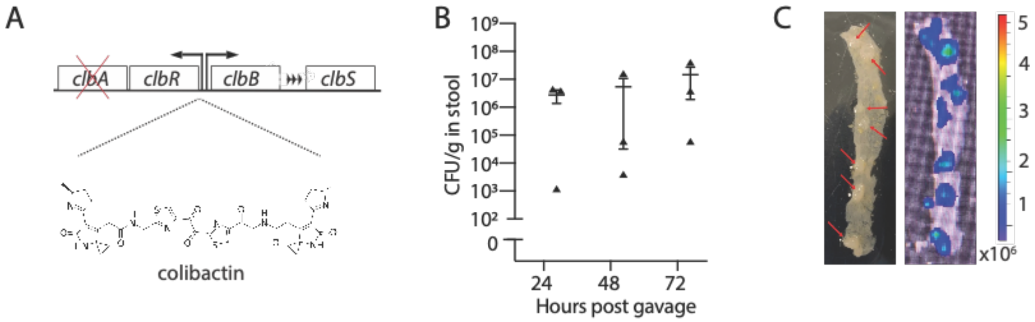


**Fig. S2. Colibactin is not a requisite for EcN colonization of intestinal adenomas. (A)** Schematic of colibactin-encoding operon in EcN whereby *clbA* is knocked out and colibactin production is disrupted. **(B-C)** 12-week-old *Apc*$^{Min/+}$ mice were gavaged twice, 3-4 days apart with $10^9$ CFU bioluminescent EcN$\Delta$*clbA*. **(C)** one stool pellet was collected 24, 48, and 72h after last dose, homogenized, plated on antibiotic-selective plates and quantified for CFU (n=3, n=1 stool per mouse). **(B)** After 1 week, mice were sacrificed, intestinal tissue was excised and *ex vivo* imaged for bioluminescence. Red arrows point to macroadenomas on distal intestinal tissue (representative image from sample size of n=5 mice).

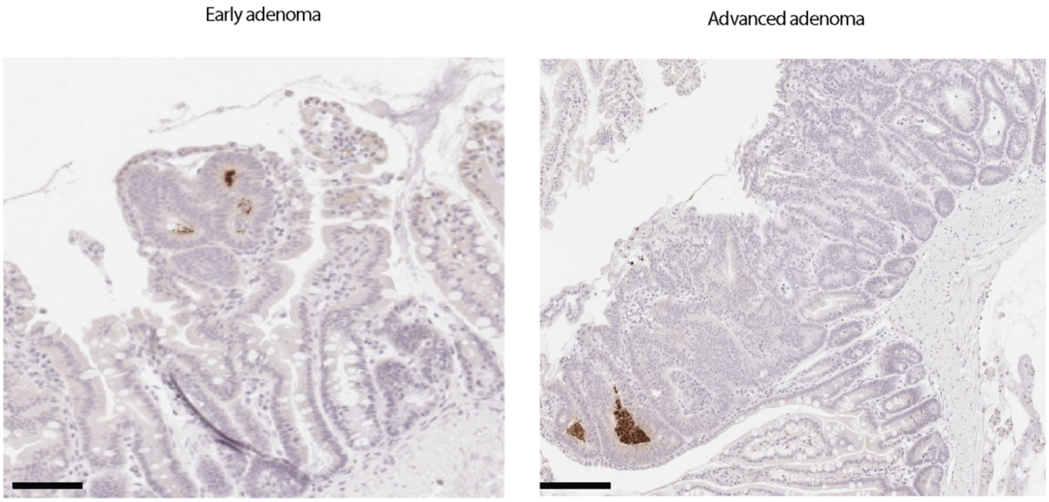


**Fig. S3. EcN can colonize and release payloads within adenomas of varying stage.** $Apc^{Min/+}$ mice were gavaged twice, 3-4 days apart, with EcN producing an HA-tagged reporter protein to enable protein detection in intestinal tissue by anti-HA immunohistochemistry after sacrifice at 4 weeks post-dosing (n=3 mice). Dark brown stain depicts HA-tagged protein in early and late-stage adenomas. Data corresponds to **Fig. 1E**.

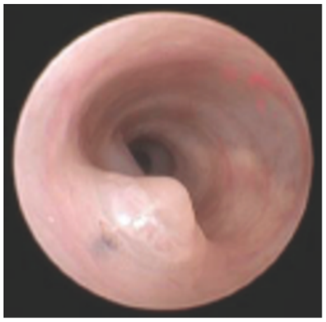

**Fig. S4. Establishment of orthotopic CRC mouse model.** Colonoscopic image of mouse tumor growing into the distal colon lumen. Orthotopic murine model of CRC was established as previously described[20].

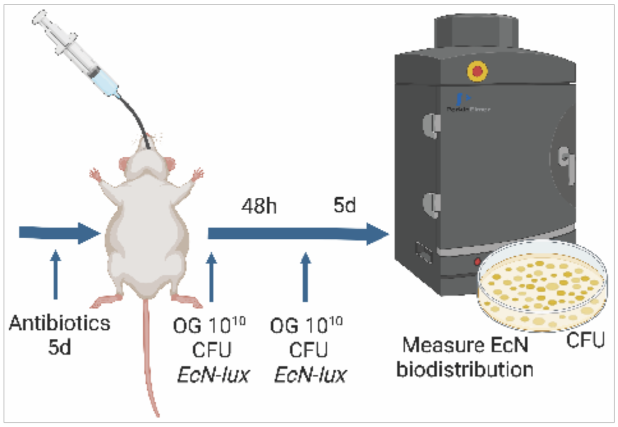


**Fig. S5.** Schematic depicting experimental timeline of data shown in **Fig. 2.**

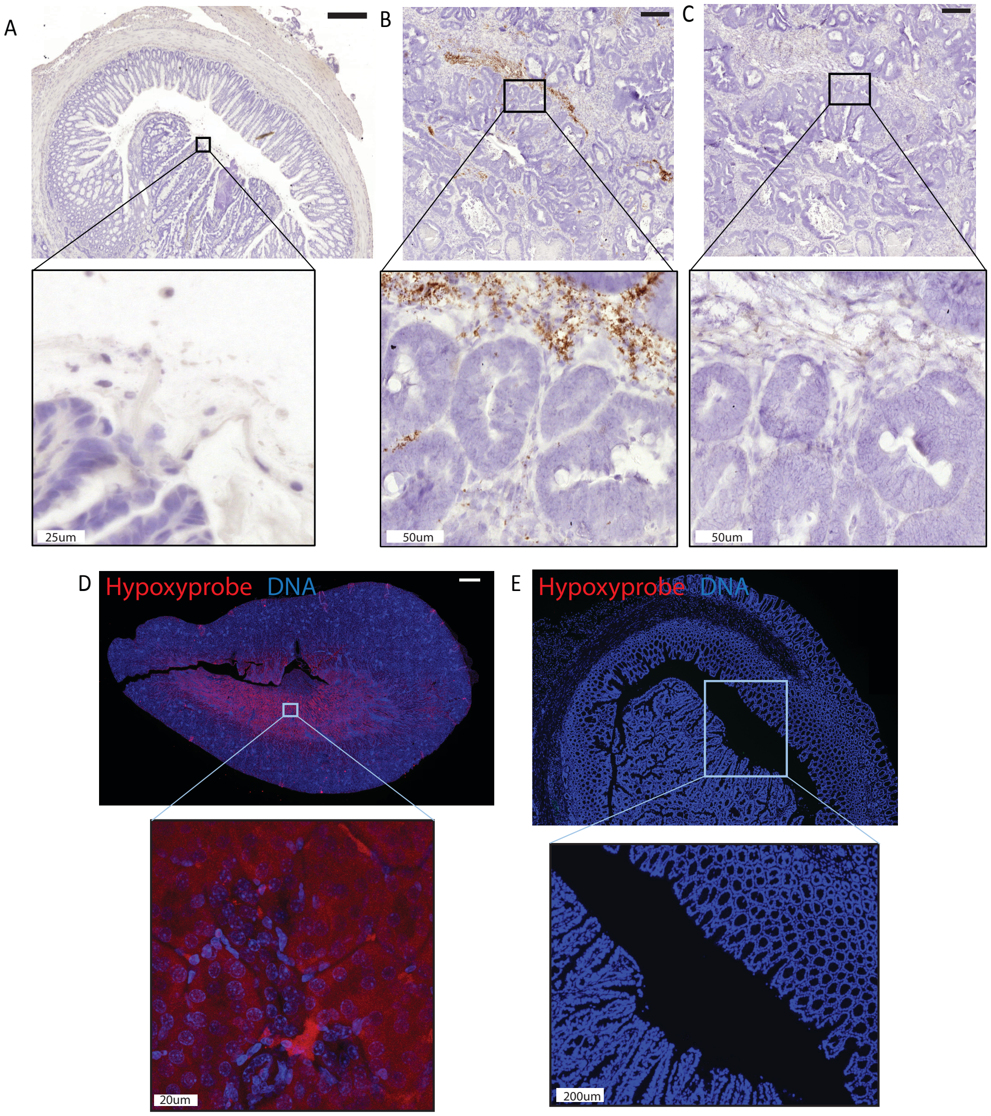


**Fig. S6. Staining for the presence of EcN in orthotopic tissues. (A)** Orthotopic CRC tumour from mouse orally-dosed twice, 2 days apart, with EcN-lux. Representative RNA ISH staining for negative control probe, *DapB* (brown) on adjacent section to Figure2F. Scale bar: 250μm & 25μm (lower inset). (**B-C)** Mouse CRC tissue intratumorally injected with EcN-lux bacteria after harvest, to generate a positive control bacteria containing tissue *ex vivo*. Representative RNAscope ISH staining for (**B)** EcN-lux (brown) and (**C)** negative control probe, *DapB* (brown). Scale bar: 200μm & 50μm (lower inset). (**D)** Representative IF staining of positive control tissue, kidney medulla, for Hypoxyprobe (red). Scale bar: 500μm & 20μm (lower inset). (**E)** Representative IF staining for Hypoxyprobe of a negative control orthotopic CRC tumour from a mouse that was injected with PBS rather than Hypoxyprobe agent prior to harvest. Scale bar: 200μm.

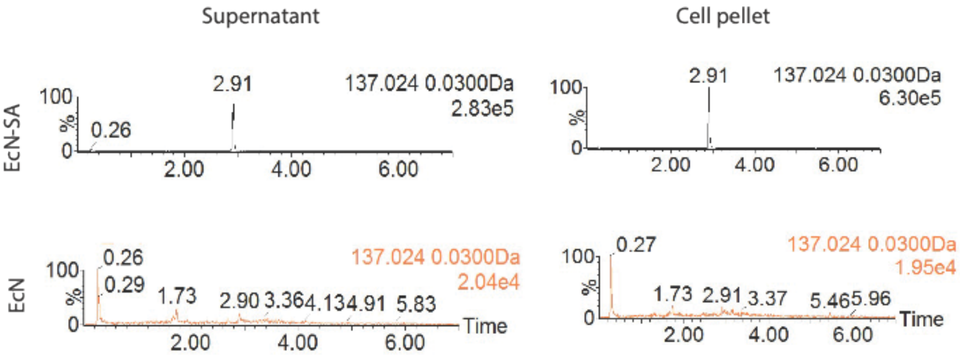


**Fig. S7. EcN can produce salicylate molecules to be detected by liquid-chromatography mass spectrometry.** EcN was engineered to produce salicylate molecules (EcN-SA). Extracted ion chromatogram showing the characteristic retention time and detected salicylate peak from EcN (negative control) and EcN-SA in cell pellets and media. Corresponds to **Fig. 3E**.

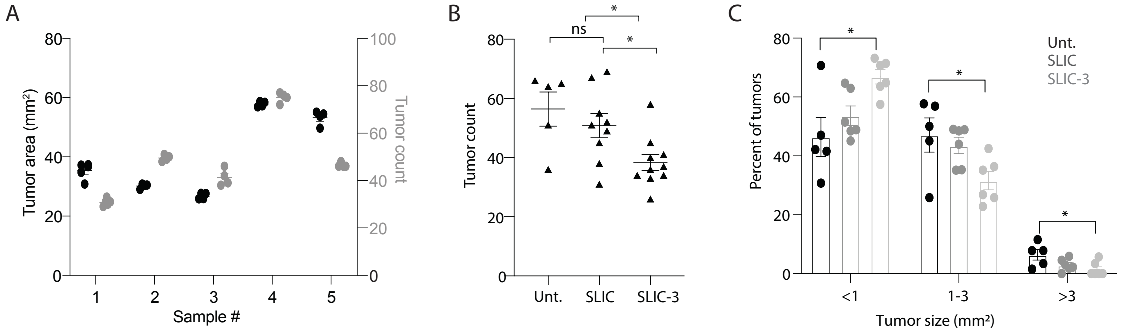


**Fig. S8. Orally-delivered EcN producing PD-L1 and CTLA-4 blocking nanobodies and GM-CSF reduces tumor burden in *Apc*$^{Min/+}$ mice.** 15–17-week-old *Apc*$^{Min/+}$ mice were dosed with PBS (Unt), EcN genomically encoding a lysis circuit (SLIC) or SLIC producing granulocyte-macrophage colony-stimulating factor (GM-CSF), and blocking nanobodies against PD-L1 and CTLA-4 targets (SLIC-3). 1 month after dosing, mice were sacrificed, intestines were bisected, swiss-rolled, paraffin embedded, sectioned, stained with hemotoxylin and eosin. Multiple sections (n=4-5 sections) in the same sample (n=5 samples) were quantified for **(A)** tumor area and tumor count to ensure that a single section was indeed representative of tumor metrics, (**B**) total tumor count based on a single section and **(C)** percent of tumors <1 mm$^2$, 1-3 mm$^2$, or <3 mm$^2$. Corresponds to data shown in **Fig. 4B**.
